# Potently neutralizing human antibodies that block SARS-CoV-2 receptor binding and protect animals

**DOI:** 10.1101/2020.05.22.111005

**Authors:** Seth J. Zost, Pavlo Gilchuk, James Brett Case, Elad Binshtein, Rita E. Chen, Joseph X. Reidy, Andrew Trivette, Rachel S. Nargi, Rachel E. Sutton, Naveenchandra Suryadevara, Lauren E. Williamson, Elaine C. Chen, Taylor Jones, Samuel Day, Luke Myers, Ahmed O. Hassan, Natasha M. Kafai, Emma S. Winkler, Julie M. Fox, James J. Steinhardt, Kuishu Ren, Yueh-Ming Loo, Nicole L. Kallewaard, David R. Martinez, Alexandra Schäfer, Lisa E. Gralinski, Ralph S. Baric, Larissa B. Thackray, Michael S. Diamond, Robert H. Carnahan, James E. Crowe

## Abstract

The COVID-19 pandemic is a major threat to global health for which there are only limited medical countermeasures, and we lack a thorough understanding of mechanisms of humoral immunity^1,2^. From a panel of monoclonal antibodies (mAbs) targeting the spike (S) glycoprotein isolated from the B cells of infected subjects, we identified several mAbs that exhibited potent neutralizing activity with IC_50_ values as low as 0.9 or 15 ng/mL in pseudovirus or wild-type (*wt*) SARS-CoV-2 neutralization tests, respectively. The most potent mAbs fully block the receptor-binding domain of S (S_RBD_) from interacting with human ACE2. Competition-binding, structural, and functional studies allowed clustering of the mAbs into defined classes recognizing distinct epitopes within major antigenic sites on the S_RBD_. Electron microscopy studies revealed that these mAbs recognize distinct conformational states of trimeric S protein. Potent neutralizing mAbs recognizing unique sites, COV2-2196 and COV2-2130, bound simultaneously to S and synergistically neutralized authentic SARS-CoV-2 virus. In two murine models of SARS-CoV-2 infection, passive transfer of either COV2-2916 or COV2-2130 alone or a combination of both mAbs protected mice from severe weight loss and reduced viral burden and inflammation in the lung. These results identify protective epitopes on the S_RBD_ and provide a structure-based framework for rational vaccine design and the selection of robust immunotherapeutic cocktails.

---

The S protein of SARS-CoV-2 is the molecular determinant of viral attachment, fusion, and entry into host cells^3^. The cryo-EM structure of a prefusion-stabilized trimeric S protein ectodomain (S2P_ecto_) for SARS-CoV-2 reveals similar features to that of the SARS-CoV S protein^4^. This type I integral membrane protein and class I fusion protein possesses an N-terminal subunit (S1) that mediates binding to receptor and a C-terminal subunit (S2) that mediates virus–cell membrane fusion. The S1 subunit contains an N-terminal domain (S_NTD_) and a receptor-binding domain (S_RBD_). SARS-CoV-2 and SARS-CoV, which share approximately 78% sequence identity in their genomes^1^ both use human angiotensin-converting enzyme 2 (hACE2) as an entry receptor^5-7^. Previous studies of human immunity to other high-pathogenicity zoonotic betacoronaviruses including SARS-CoV^8-12^ and Middle East respiratory syndrome (MERS)^13-22^ showed that Abs to the viral surface spike (S) glycoprotein mediate protective immunity. The most potent S protein-specific mAbs appear to neutralize betacoronaviruses by blocking attachment of virus to host cells by binding to the region on S_RBD_ that directly mediates engagement of the receptor. It is likely that human Abs have promise for use in modifying disease during SARS-CoV-2 infection, when used for prophylaxis, post-exposure prophylaxis, or treatment of SARS-CoV-2 infection^23^. Many studies including randomized controlled trials evaluating convalescent plasma and one trial evaluating hyperimmune immunoglobulin are ongoing, but it is not yet clear whether such treatments can reduce morbidity or mortality^24^.

We isolated a large panel of SARS-CoV-2 S protein-reactive mAbs from the B cells of two individuals who were previously infected with SARS-CoV-2 in Wuhan China^25^. A subset of those antibodies bound to the receptor-binding domain of S (S_RBD_) and exhibited neutralizing activity in a rapid screening assay with authentic SARS-CoV-2^25^. Here, we defined the antigenic landscape of SARS-CoV-2 and determined which sites of S_RBD_ are the target of protective mAbs. We tested a panel of 40 anti-S human mAbs we previously pre-selected by a rapid neutralization screening assay in a quantitative focus reduction neutralization test (FRNT) with SARS-CoV-2 strain WA1/2020. These assays revealed the panel exhibited a range of half-maximal inhibitory concentration (IC_50_) values, from 15 to over 4,000 ng/mL (visualized as a heatmap in Fig. 1a, values shown in Extended Data Table 1, and full curves shown in Extended Data Fig. 1). We hypothesized that many of these S_RBD_-reactive mAbs neutralize virus infection by blocking S_RBD_ binding to hACE2. Indeed, most neutralizing mAbs we tested inhibited the interaction of hACE2 with trimeric S protein directly (Fig. 1a, Extended Data Fig. 2). Consistent with these results, these mAbs also bound strongly to a trimeric S ectodomain (S2P_ecto_) protein or monomeric S_RBD_ (Fig. 1a, Extended Data Fig. 3). We evaluated whether S2P_ecto_ or S_RBD_ binding or hACE2-blocking potency predicted binding neutralization potency independently, but none of these measurements correlated with neutralization potency (Fig. 1b-d). However, each of the mAbs in the highest neutralizing potency tier (IC_50_<150 ng/mL) also revealed strongest blocking activity against hACE2 (IC_50_< 150 ng/mL) and exceptional binding activity (EC_50_< 2 ng/mL) to S2P_ecto_ trimer and S_RBD_ (Fig. 1e). Representative neutralization curves for two potently neutralizing mAbs designated COV2-2196 and COV2-2130 are shown in (Fig. 1f). Potent neutralization was confirmed using pseudovirus neutralization assays, which revealed far more sensitive neutralization phenotypes than *wt* virus and demonstrated a requirement for the use of live virus assays for assessment of mAb potency (Fig. 1g). Both of these mAbs bound strongly to S2P_ecto_ trimer and fully blocked hACE2 binding (Fig. 1h-i).

**Figure 1.**
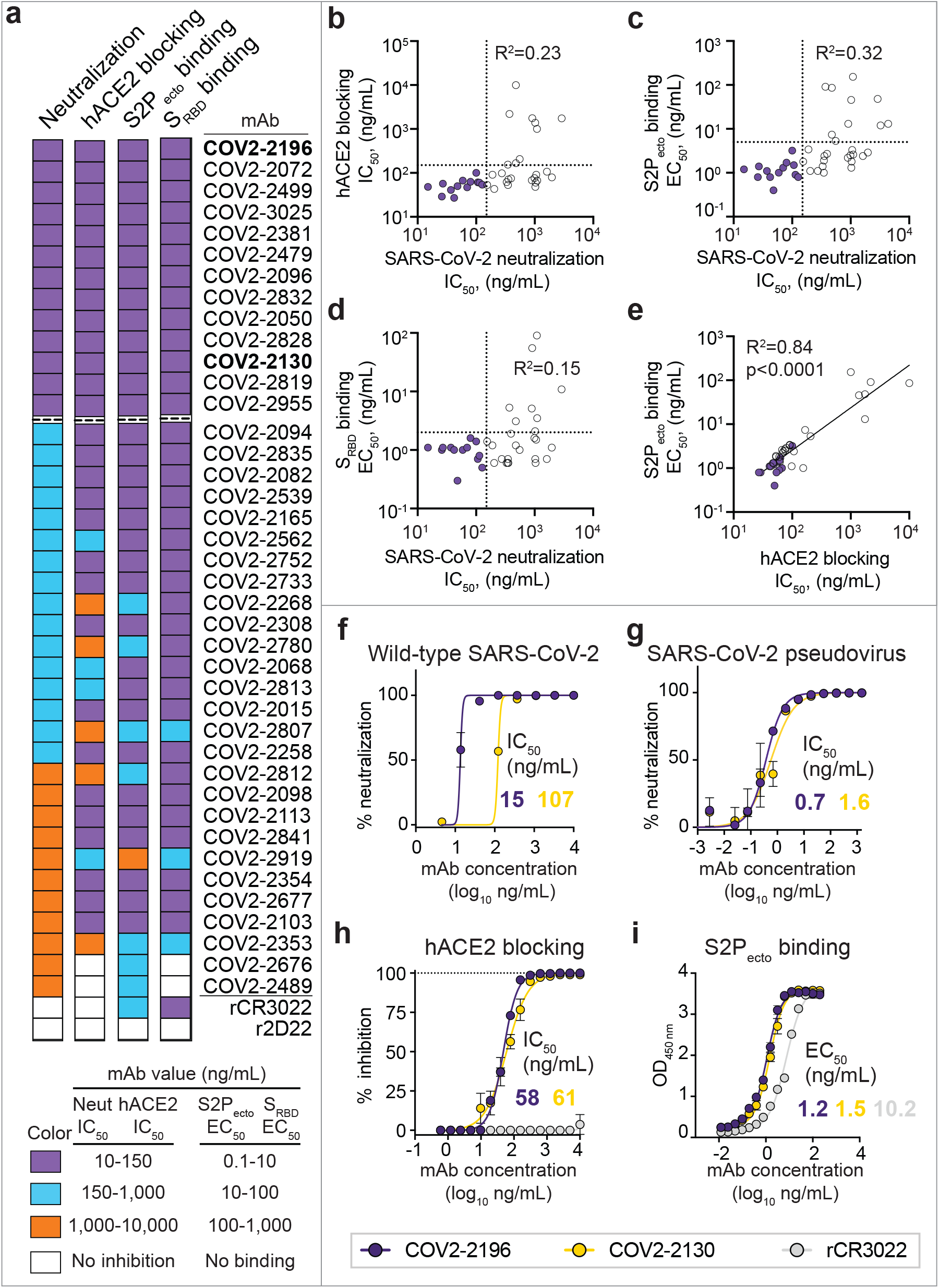
Functional characteristics of neutralizing SARS-CoV-2 mAbs. **a.** Heatmap of mAb neutralization activity, hACE2 blocking activity, and binding to either trimeric S2Pecto protein or monomeric SRBD. MAbs are ordered by neutralization potency (highest at the top, lowest at the bottom). Dashed lines indicate the 13 antibodies with a neutralization IC50 value lower than 150 ng/mL for wt virus. IC50 values are visualized for viral neutralization and hACE2 blocking, while EC50 values are visualized for binding. A recombinant form of the cross-reactive SARS-CoV SRBD mAb CR3022 is shown as a positive control, while the anti-dengue mAb 2D22 is shown as a negative control. Data are representative of at least 2 independent experiments, each performed in technical duplicate. No inhibition indicates an IC50 value of >10,000 ng/mL, while no binding indicates an EC50 value of >10,000 ng/mL. **b-d.** Correlation of hACE2 blocking, S2Pecto trimer binding, or SRBD binding of mAbs with their neutralization activity. R2 values are shown for linear regression analysis of log-transformed values. Purple circles indicate mAbs with a neutralization IC50 value lower than 150 ng/mL. **e.** Correlation of hACE2 blocking and S2Pecto trimer binding. R2 values are shown for linear regression analysis of log-transformed values. **f.** Neutralization curves for COV2-2196 and COV2-2130 in a neutralization assay against authentic SARS-CoV-2 virus. Calculated IC50 values are denoted on the graph. Error bars denote the standard deviation of each point. Data are representative of at least 2 independent experiments, each performed in technical duplicate. **g.** Neutralization curves for COV2-2196 and COV2-2130 in a pseudovirus neutralization assay. Error bars denote the standard deviation of each point. Values shown are technical duplicates from a single experiment. Calculated IC50 values from a minimum of 6 experiments are denoted on the graph. **h.** hACE2 blocking curves for COV2-2196, COV2-2130, and the non-blocking SARS-CoV mAb rCR3022 in the hACE2 blocking ELISA. Calculated IC50 values are denoted on the graph. Error bars denote the standard deviation of each point. Values shown are technical triplicates from a representative experiment repeated twice. **i.** ELISA binding of COV2-2196, COV2-2130, and rCR3022 to trimeric S2Pecto. Calculated EC50 values are denoted on the graph. Error bars denote the standard deviation of each point. Values shown are technical triplicates from a representative experiment repeated twice.

We next defined the major antigenic sites on S_RBD_ for neutralizing mAbs by competition-binding analysis. We first used a biolayer interferometry-based competition assay with a minimal S_RBD_ domain to screen for mAbs that competed for binding with the potently neutralizing mAb COV2-2196 or a recombinant version of the previously described SARS-CoV mAb CR3022, which recognizes a conserved cryptic epitope^10,26^. We identified three major groups of competing mAbs (Fig. 2a). The largest group of mAbs blocked COV2-2196 but not rCR3022, while some mAbs were blocked by rCR3022 but not COV2-2196. Two mAbs, including COV2-2130, were not blocked by either reference mAb. Most mAbs competed with hACE2 for binding, suggesting that they bound near the hACE2 binding site of the S_RBD_. We used COV2-2196, COV2-2130, and rCR3022 in an ELISA-based competition-binding assay with trimeric S2P_ecto_ protein and also found that S_RBD_ contained three major antigenic sites, with some mAbs likely making contacts in more than one site (Fig. 2b). Most of the potently neutralizing mAbs directly competed for binding with COV2-2196.

**Figure 2.**
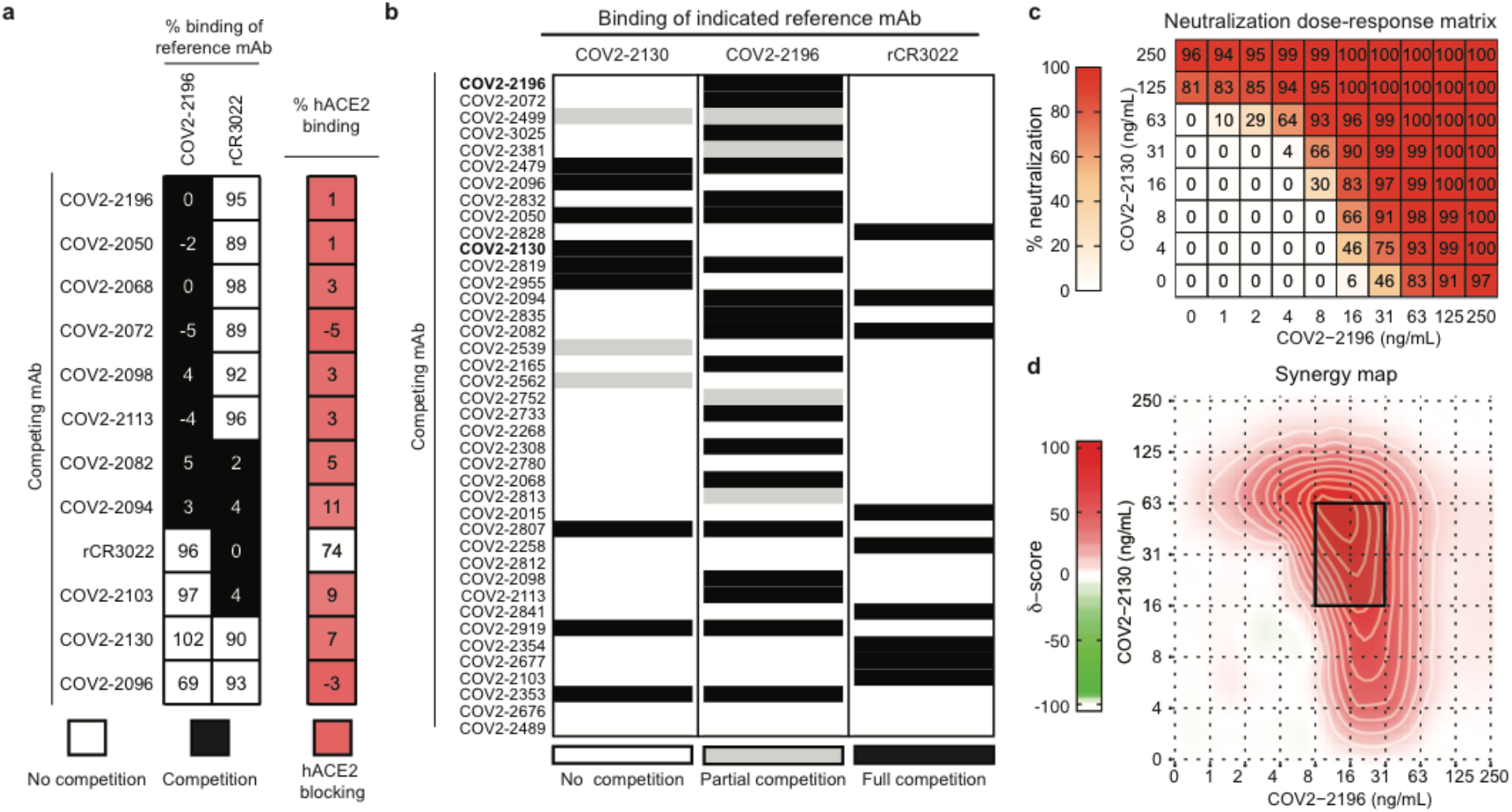
Epitope mapping of mAbs by competition-binding analysis and synergistic neutralization by a pair of mAbs. **a.** Left: biolayer interferometry-based competition binding assay measuring the ability of mAbs to prevent binding of reference mAbs COV2-2196 and rCR3022 to RBD fused to mouse Fc (RBD-mFc) loaded onto anti-mouse Fc biosensors. Values in squares are % of binding of the reference mAb in the presence of the competing mAb relative to a mock-competition control. Black squares denote full competition (<33% of binding relative to no-competition control), while white squares denote no competition (>67% of binding relative to no-competition control). Right: biolayer interferometry-based competition binding assay measuring the ability of mAbs to prevent binding of hACE2. Values denote % binding of hACE2, normalized to hACE2 binding in the absence of competition. Red color denotes competition of mAb with hACE2. **b.** Competition of neutralizing mAb panel with reference mAbs COV2-2130, COV2-2196, or rCR3022. Reference mAbs were biotinylated and binding of reference mAbs to trimeric S2Pecto was measured in the presence of saturating amounts of each mAb in a competition ELISA. ELISA signal for each reference mAb was normalized to the signal in the presence of the non-binding anti-dengue mAb 2D22. Black denotes full competition (<25% binding of reference mAb), grey denotes partial competition (25-60% binding of reference mAb), and white denotes no competition (>60% binding of reference mAb). **c.** Neutralization dose-response matrix of wild-type SARS-CoV-2 by COV2-2196 and COV2-2130. Axes denote the concentration of each mAb. Experiment was performed in technical triplicate. Shown is a representative experiment that was performed in technical triplicate. % neutralization for each combination of mAbs is shown in each square. A white-to-red heatmap denotes 0% neutralization to 100% neutralization, respectively. **d.** Synergy map calculated based on the SARS-CoV-2 neutralization in the above panel. Red color denotes areas where synergistic neutralization was observed, and a black box denotes the area of maximal synergy between the two mAbs.

Since COV2-2196 and COV2-2130 did not compete for binding to S_RBD_, we assessed if these mAbs synergize for virus neutralization, a phenomenon previously observed for SARS-CoV mAbs^10^. We tested combination responses (see dose-response neutralization matrix, Fig. 2c) in the FRNT using SARS-CoV-2 and compared these experimental values with the expected responses calculated by synergy scoring models^27^. The comparison revealed that the combination of COV2-2196 + COV2-2130 was synergistic (with a synergy score of 17.4, where any score of >10 indicates synergy). The data in Fig. 2c shows the dose-response synergy matrix and demonstrates that a combined mAb dose of 79 ng/mL in the cocktail (16 ng/mL of COV2-2196 and 63 ng/mL of COV2-2130) had the same activity as 250 ng/mL of each individual mAb (see Fig. 2c). This finding shows that using a cocktail the dose of each mAb can be reduced by more than three-fold to achieve the same potency of virus neutralization *in vitro*.

We next defined the epitopes recognized by representative mAbs in the two major competition-binding groups that synergize for neutralization. We performed mutagenesis studies of the S_RBD_ using alanine or arginine substitution to determine critical residues for binding of neutralizing mAbs (Extended Data Fig. 4). Loss of binding studies revealed F486A or N487A as critical residues for COV2-2196 and N487A as a critical residue for COV2-2165, which compete with one another for binding, and likewise mutagenesis studies for COV2-2130 using K444A and G447R mutants defined these residues as critical for recognition (Fig. 3a). Previous structural studies have defined the interaction between the S_RBD_ and hACE2 (Fig. 3b)^28^. Most of the interacting residues in the S_RBD_ are contained within a 60-amino-acid linear peptide that defines the hACE2 recognition motif (Fig. 3c). We next tested binding of human mAbs to this minimal peptide and found that potent neutralizing members of the largest competition-binding group including COV2-2196, COV2-2165, and COV2-2832 recognized this peptide (Fig. 3c), suggesting these mAbs make critical contacts within the hACE2 recognition motif.

**Figure 3.**
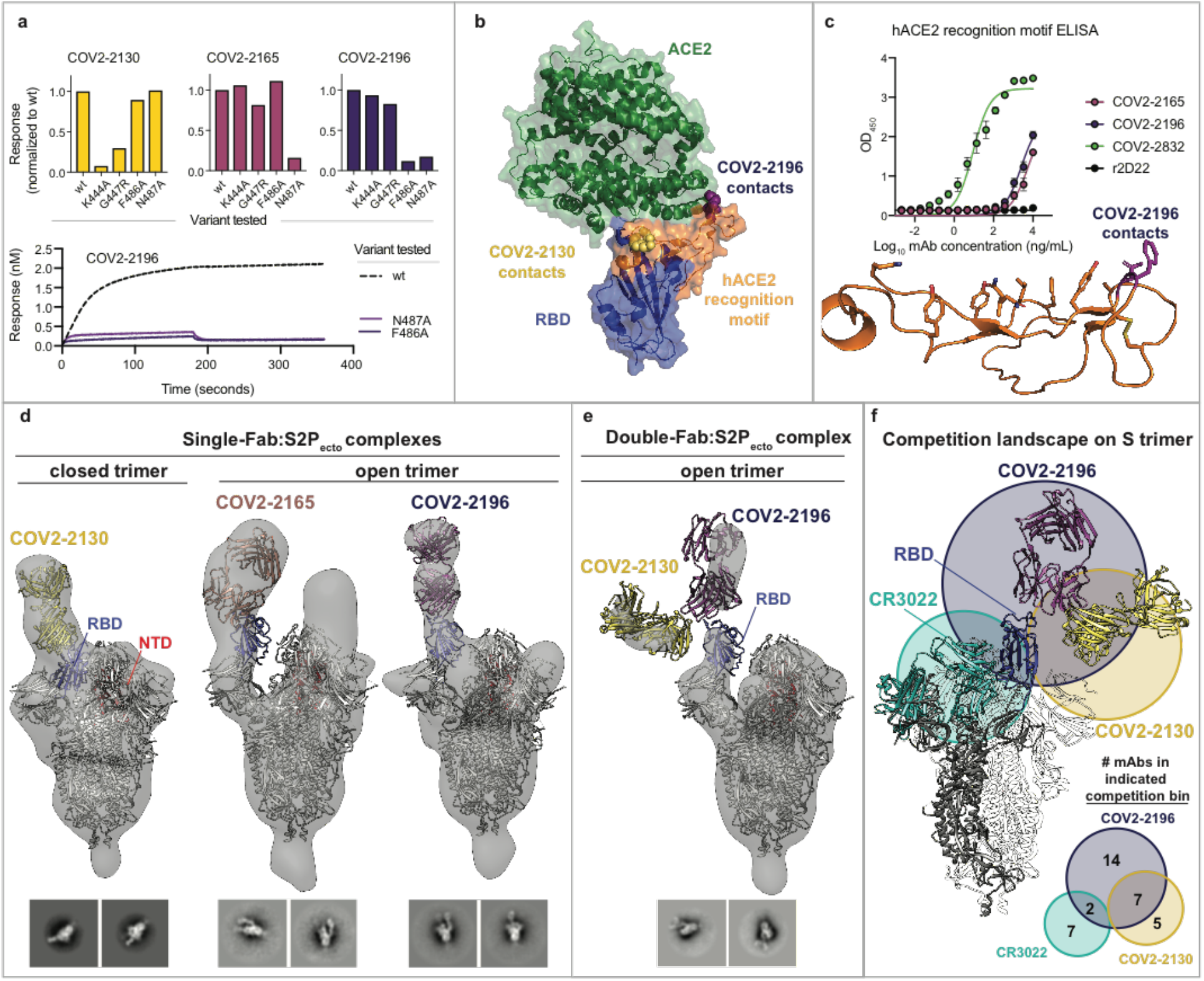
Epitope identification and structural characterization of mAbs. **a.** Identification of critical contact residues by alanine and arginine mutagenesis. Top: binding of COV2-2130 (gold), COV2-2165 (maroon) or COV2-2196 (dark purple) to wild-type (wt) or mutant SRBD constructs measured by biolayer interferometry. Shown on y-axis is the response normalized to the signal observed for binding to wt SRBD. Bottom: representative binding curves for COV2-2196 to wt or SRBD constructs with critical contact residues mutated. **b.** Crystal structure of SARS-CoV-2 (blue) and hACE2 (green) (PDB (6M0J). The hACE2 recognition motif is colored orange. Critical contact residues for COV2-2130 are shown as gold spheres, while critical contact residues for COV2-2196 are shown as purple spheres. **c.** ELISA binding of mAbs to the 60-amino-acid hACE2 recognition motif. r2D22, an anti-dengue mAb, is shown as a negative control. Bottom: structure of hACE2 recognition motif in orange with COV2-2196 critical contact residues shown in purple. **d.** Single-Fab:S2Pecto trimer complexes visualized by negative-stain electron microscopy for COV2-2130 (gold), COV2-2165 (maroon), or COV2-2196 (dark purple). The RBD is shown in blue and the S N-terminal domain (NTD) is shown in red. Electron density is shown in grey. Trimer state (open or closed) is denoted for each complex. Representative 2D class averages for each complex are shown at the bottom (box size 128 pixel). **e.** COV2-2130 and COV2-2196 Fabs in complex with S2Pecto trimer. Simultaneous binding of COV2-2130 (gold) and COV2-2196 (purple) Fabs to S2Pecto trimer. Electron density is shown in grey. Trimer state (open or closed) is denoted. Representative 2D class averages for the complexes are shown at the bottom (box size 128 pixels). All images were made with Chimera. Competition-binding analysis visualized on S2Pecto trimer. The CR3022 crystal structure was docked into the double-Fab:S2Pecto trimer structure. CR3022 is shown in cyan. Bottom: a quantitative Venn diagram notes the number of mAbs in each competition group and the overlap between groups.

We used negative-stain electron microscopy of S2P_ecto_ trimer/Fab complexes to structurally determine the epitopes for these mAbs. The potently neutralizing antibodies COV2-2196 and COV2-2165 bound to the hACE2 recognition motif of S_RBD_ and recognized the ‘open’ conformational state of the S2P_ecto_ trimer (Fig. 3d). The mode of engagement of these two antibodies differed, however, as the binding pose and the angle relative to the spike ‘body’ for one was different compared to the other. COV2-2130, which represents the second competition-binding group, bound to the RBD in the S2P_ecto_ trimer in the ‘closed’ position (Fig. 3d). Since COV2-2196 and COV2-2130 did not compete for binding, we attempted to make complexes of both Fabs bound at the same time to the S2P_ecto_ trimer. We found that both Fabs bound simultaneously when the S2P_ecto_ trimer was in the open position, indicating that COV2-2130 can recognize the S_RBD_ in both conformations (Fig. 3e). Overlaying the two-Fab complex with the structure of the RBD:CR3022 complex^26^, we observed that these antibodies bind to three distinct sites on S_RBD_, as predicted based on our competition-binding studies (Fig. 3f).

Next, we tested the prophylactic efficacy of COV2-2196 or COV2-2130 monotherapy or a combination of COV2-2196 + COV2-2130 in a newly developed SARS-CoV-2 infection model in BALB/c mice in which hACE2 is expressed in the lung after intranasal adenovirus (AdV-hACE2) transduction. In this relatively stringent disease model, we also administered a single dose of anti-Ifnar1 antibody to augment virus infection and pathogenesis, which results in a disseminated interstitial pneumonia (A. Hassan and M. Diamond, submitted for publication). We passively transferred a single dose of mAb COV2-2196 (10 mg/kg), COV2-2130 (10 mg/kg), a combination of COV2-2196 + COV2-2130 (5 mg/kg each), or an isotype control mAb (10 mg/kg) to AdV-hACE2-transduced mice one day before intranasal challenge with 4 × 10^5^ PFU of SARS-CoV-2. Prophylaxis with COV2-2196 or COV2-2130 or their combination prevented severe SARS-CoV-2-induced weight loss through the first week of infection (Fig. 4a). Viral RNA levels were reduced significantly at 7 dpi in the lung and distant sites including the heart and spleen (Fig. 4b). The expression of interferon gamma (INF-g), IL-6, CXCL10 and CCL2 cytokine and chemokine genes, which are indicators of inflammation, also was reduced in the lung of treated mice at 7 dpi —the peak of the disease (Fig. 4c).

**Figure 4.**
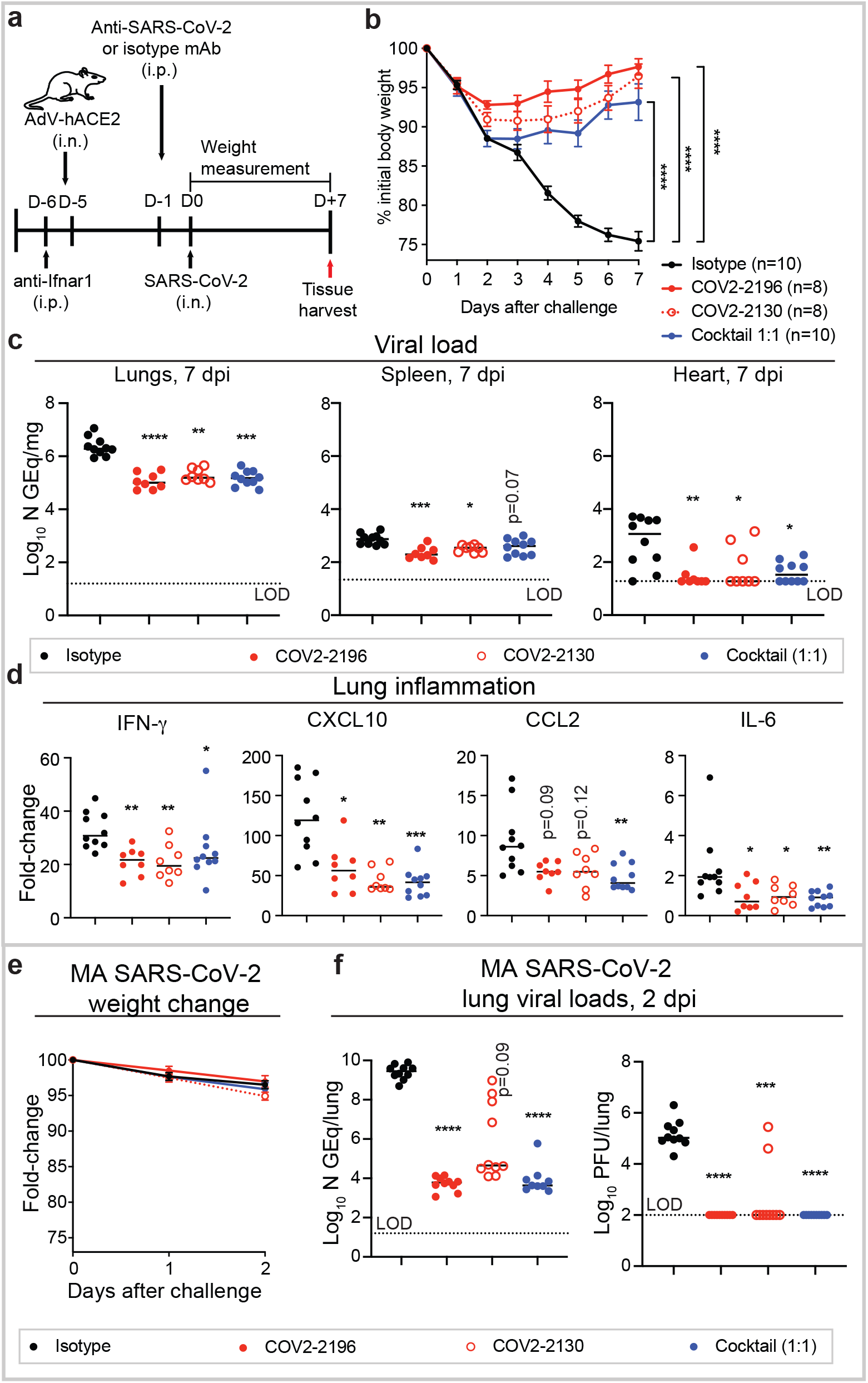
Protective efficacy of neutralizing human mAbs against SARS-CoV-2 infection. **a.** SARS-CoV-2 challenge model. Ten to eleven-week-old BALB/c mice (two experiments of 4-5 mice per group) were treated with anti-Ifnar1 mAb and transduced with AdV-hACE2 via the i.n. route one day later. After four days, mice were treated via the i.p. route with 200 μg of mAbs CoV2-2196, -2130, or combination (1:1 ratio) or isotype control mAb. One day later, SARS-CoV-2 was inoculated via the i.n. route. Tissues were harvested at 7 dpi for analysis (c, d). **b.** Body weight change of mice in panel a. (two-way ordinary ANOVA with Tukey’s post-test: **** P < 0.0001). **c.** Viral burden in the lung, spleen and heart was measured by RT-qPCR: Kruskal-Wallis ANOVA with Dunn’s post-test (*, P < 0.05, ** P < 0.01, *** P < 0.001, **** P < 0.0001). The dashed line indicates the assay limit of detection. **d.** Cytokine and chemokine gene expression was measured by qPCR analysis. Kruskal-Wallis ANOVA with Dunn’s post-test (*, P < 0.05, ** P < 0.01, *** P < 0.001). **e.** MA-SARS-CoV-2 challenge model. Twelve-week-old BALB/c mice (n=10) were inoculated with 105 PFU of MA-SARS-CoV-2 via the i.n. route. Body weight change of mice is shown. **f.** Viral burden in the lung was measured at 2 dpi by RT-qPCR (left) or plaque assay (right) from (e): Kruskal-Wallis ANOVA with Dunn’s post-test (*** P < 0.001, **** P < 0.0001).

We also tested COV2-2196 or COV2-2130 or their combination for prophylactic efficacy in an immunocompetent model using a mouse-adapted (MA) SARS-CoV-2 virus^29^ (Fig. 4d). *In vitro* tests showed that the IC_50_ values for neutralization were comparable for the *wt* and MA SARS-CoV-2 viruses for these mAbs (data not shown). Each of the mAb treatments delivered at a dose of 200 μg/mouse (~ 8 mg/kg) reduced viral RNA levels up to 10^5^-fold at 2 dpi in the lung when compared to the isotype control group (Fig. 4e, left). Concordantly, all animals from COV2-2196 and COV2-2196 + COV2-2130 treatment group and 8 of 10 animals from COV2-2130 treatment no longer had infectious virus at 2 dpi in the lung as measured by plaque titer of lung tissue (Fig. 4e, right). Collectively, these results in mice suggested that COV2-2196 or COV2-2130 alone or in combination are promising candidates for treatment or prevention of COVID-19.

Here, we defined the antigenic landscape for a large panel of highly potent mAbs against SARS-CoV-2. These detailed studies and the screening studies that identified this panel of mAbs from a larger panel of hundreds^25^ demonstrate that although diverse human neutralizing antibodies are elicited by natural infection with SARS-CoV-2, only a small subset of those mAbs are of high potency (IC_50_<50 ng/mL against live SARS-CoV-2 virus), and therefore, suitable for therapeutic development. Biochemical and structural analysis of these potent mAbs defined three principal antigenic sites of vulnerability to neutralization by human mAbs elicited by natural infection with SARS-CoV on the S_RBD_. Representative mAbs from the two most potent antigenic sites were shown to synergize *in vitro* and protect as an *in vivo* cocktail. This finding reveals critical features of effective humoral immunity to SARS-CoV-2 and suggests that the role of synergistic neutralization activity in polyclonal responses should be explored further. Moreover, as SARS-CoV-2 continues to circulate, population immunity elicited by natural infection may start to select for antigenic variants that escape from the selective pressure of neutralizing antibodies, reinforcing the need to target multiple epitopes of S protein in vaccines or immunotherapeutics.

The common S gene variants across the globe reported to date are located at positions D614G, V483A, L5F, Q675H, H655Y and S939F^30^, far away from the amino acid variants at residues 486 or 487 identified in our mutation studies for the lead mAbs studied here. Rationally-selected therapeutic cocktails like the one described here might offer even greater resistance to SARS-CoV-2 escape. These studies set the stage for preclinical evaluation and development of the identified mAbs as candidates for use as COVID-19 immunotherapeutics in humans.

## Data availability

The EM maps have been deposited at the Electron Microscopy Data Bank with accession codes EMBD 21965 (S2P_ecto_ apo), EMD-21974 (S2P_ecto_ + Fab COVs-2165), EMD-21975 (S2P_ecto_ + Fab COVs-2196), EMD-21976 (S2P_ecto_ + Fab COVs-2130) and EMD-21977 (S2P_ecto_ + Fab COV2-2196 + Fab COV2-2130). Materials reported in this study will be made available but may require execution of a Materials Transfer Agreement.

## Acknowledgements

We thank Angela Jones and the staff of the Vanderbilt VANTAGE core laboratory for expedited sequencing, Ross Trosseth for assistance with data management and analysis, Robin Bombardi and Cinque Soto of VUMC for technical consultation on genomics approaches, Arthur Kim, Adam Bailey, Laura VanBlargan, James Earnest, Broc McCune and Swathi Shrihari of WUSTL for experimental assistance and key reagents, and Kevin M. Tuffy, Seme Diallo, Patrick M. McTamney, and Lori Clarke of AstraZeneca for generation of protein and pseudovirus reagents and related data. This study was supported by Defense Advanced Research Projects Agency (DARPA) grants HR0011-18-2-0001 and HR00 11-18-3-0001, NIH contracts 75N93019C00074 and 75N93019C00062 and the Dolly Parton COVID-19 Research Fund at Vanderbilt. This work was supported by NIH grant 1S10RR028106-01A1 for the Next Generation Nucleic Acid Sequencer, housed in Vanderbilt Technologies for Advanced Genomics (VANTAGE) and the Vanderbilt Institute for Clinical and Translational Research with grant support from (UL1TR002243 from NCATS/NIH). S.J.Z. was supported by NIH T32 AI095202. J.B.C. is supported by a Helen Hay Whitney Foundation postdoctoral fellowship. D.R.M. was supported by NIH T32 AI007151 and a Burroughs Wellcome Fund Postdoctoral Enrichment Program Award. J.E.C. is the recipient of the 2019 Future Insight Prize from Merck KGaA, Darmstadt Germany, which supported this research with a research grant. The content is solely the responsibility of the authors and does not necessarily represent the official views of the U.S. government or the other sponsors.

## Author contributions

Conceived of the project: S.J.Z., P.G., R.H.C., L.B.T., M.S.D., J.E.C.; Obtained funding: J.E.C. and M.S.D. Performed laboratory experiments: S.J.Z., P.G., J.B.C., E.B., R.E.C., J.X.R., A.T., R.S.N., R.E.S., N.S., L.E.W., A.O.H., N.M.K., E.W., J.M.F., L.B.T., J.J.S., K.R., Y.-M.L., A.S., L.E.G., D.R.M.; Performed computational work: E.C.C., T.J., S.D., L.M.; Supervised research: N.L.K, M.S.D., L.B.T., R.S.B., R.H.C., J.E.C. Wrote the first draft of the paper: S.J.Z., P.G., R.H.C., J.E.C.; All authors reviewed and approved the final manuscript.

## Competing interests

R.S.B. has served as a consultant for Takeda and Sanofi Pasteur on issues related to vaccines. M.S.D. is a consultant for Inbios, Vir Biotechnology, NGM Biopharmaceuticals, Eli Lilly, and is on the Scientific Advisory Board of Moderna, a past recipient of unrelated research grant from Moderna and a current recipient of an unrelated research grant Emergent BioSolutions. J.E.C. has served as a consultant for Sanofi and is on the Scientific Advisory Boards of CompuVax and Meissa Vaccines, is a recipient of previous unrelated research grants from Moderna and Sanofi and is Founder of IDBiologics, Inc. Vanderbilt University has applied for patents concerning SARS-CoV-2 antibodies that are related to this work. AstraZeneca has filed patents for materials/findings related to this work. J.J.S., K.R., Y.-M.L., and N.L.K. are employees of AstraZeneca and currently hold AstraZeneca stock or stock options. All other authors declared no competing interests.

## Additional information

**Supplementary information** is available for this paper.

**Correspondence and requests for materials** should be addressed to J.E.C.

## Supplementary Information

**Extended Data Figure. 1.**
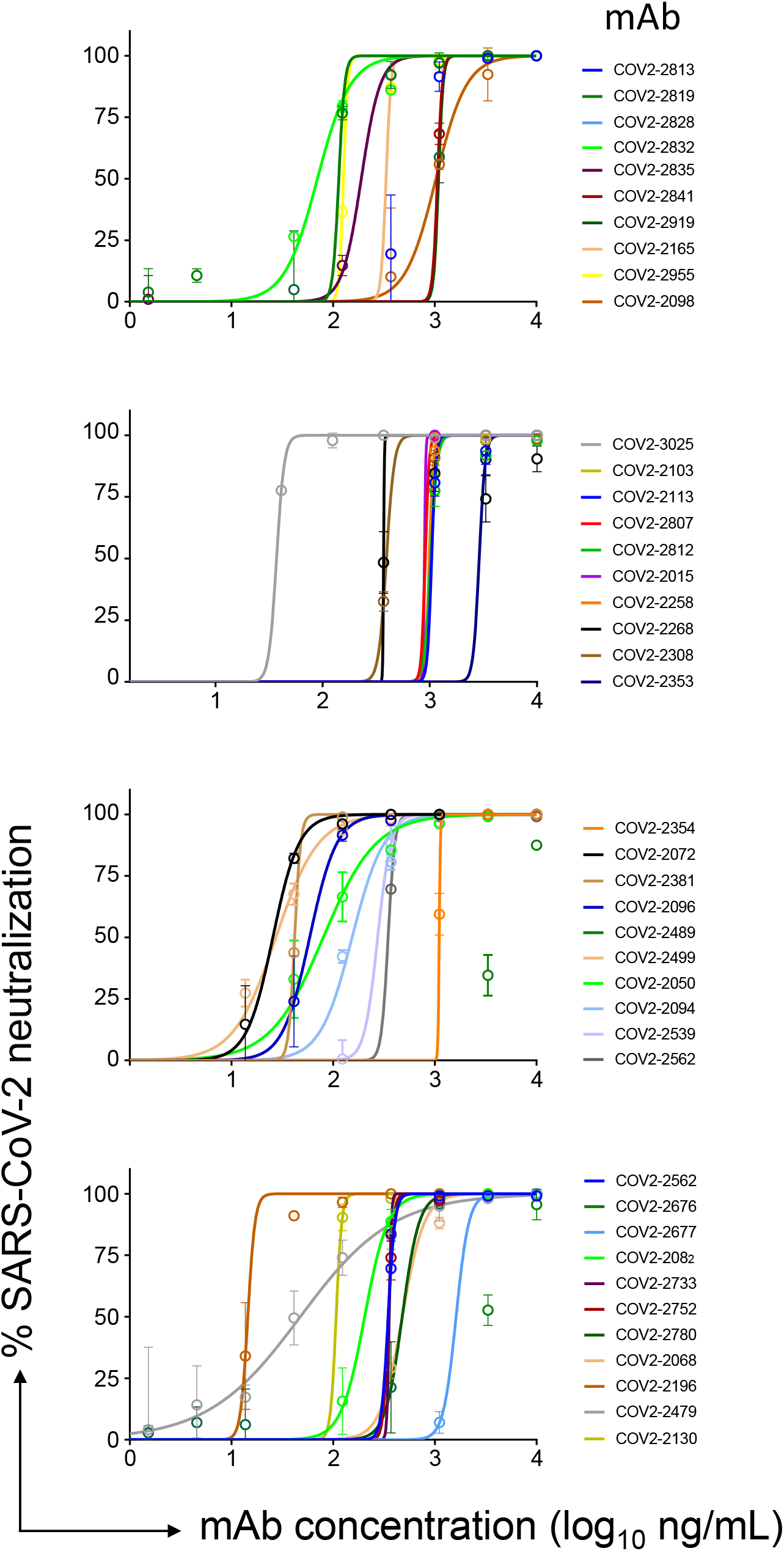
SARS-CoV-2 neutralization curves for mAb panel. Neutralization of authentic SARS-CoV-2 by human mAbs. Mean ± SD of technical duplicates is shown. Data represent one of two or more independent experiments

**Extended Data Figure 2.**
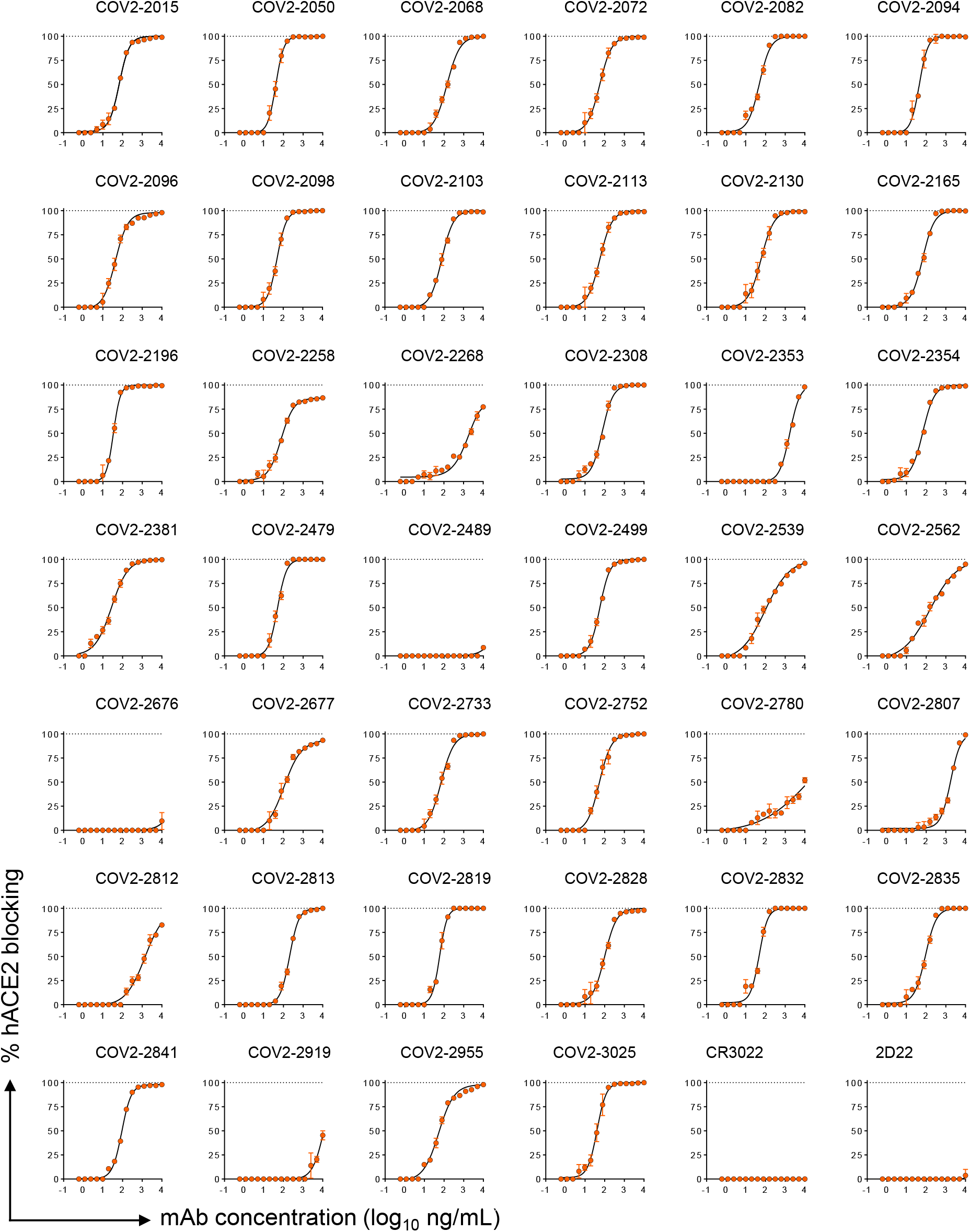
Inhibition curves for mAb inhibition of S2P_ecto_ binding to hACE2. Blocking of hACE2 binding to S2P_ecto_ by anti-SARS-CoV-2 neutralizing human mAbs. Mean ± SD of triplicates of one experiment is shown. Antibodies CR3022 and 2D22 served as controls.

**Extended Data Figure. 3.**
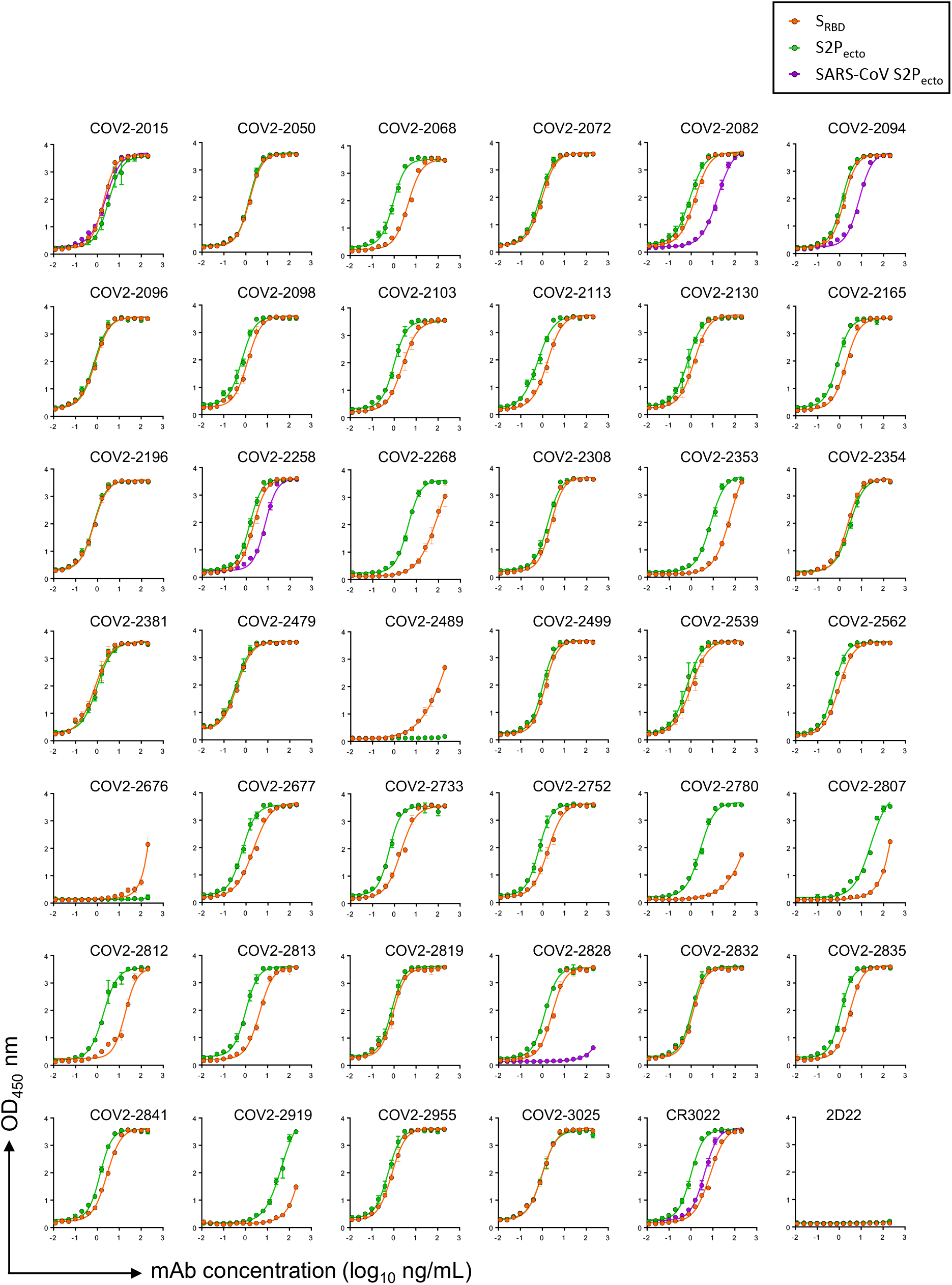
ELISA binding of anti-SARS-CoV-2 neutralizing human mAbs to trimeric S_RBD_, S2P_ecto_, or SARS-CoV S2P_ecto_ antigen. Mean ± SD of triplicates and representative of two experiments are shown. Antibodies CR3022 and 2D22 served as controls.

**Extended Data Figure. 4.**
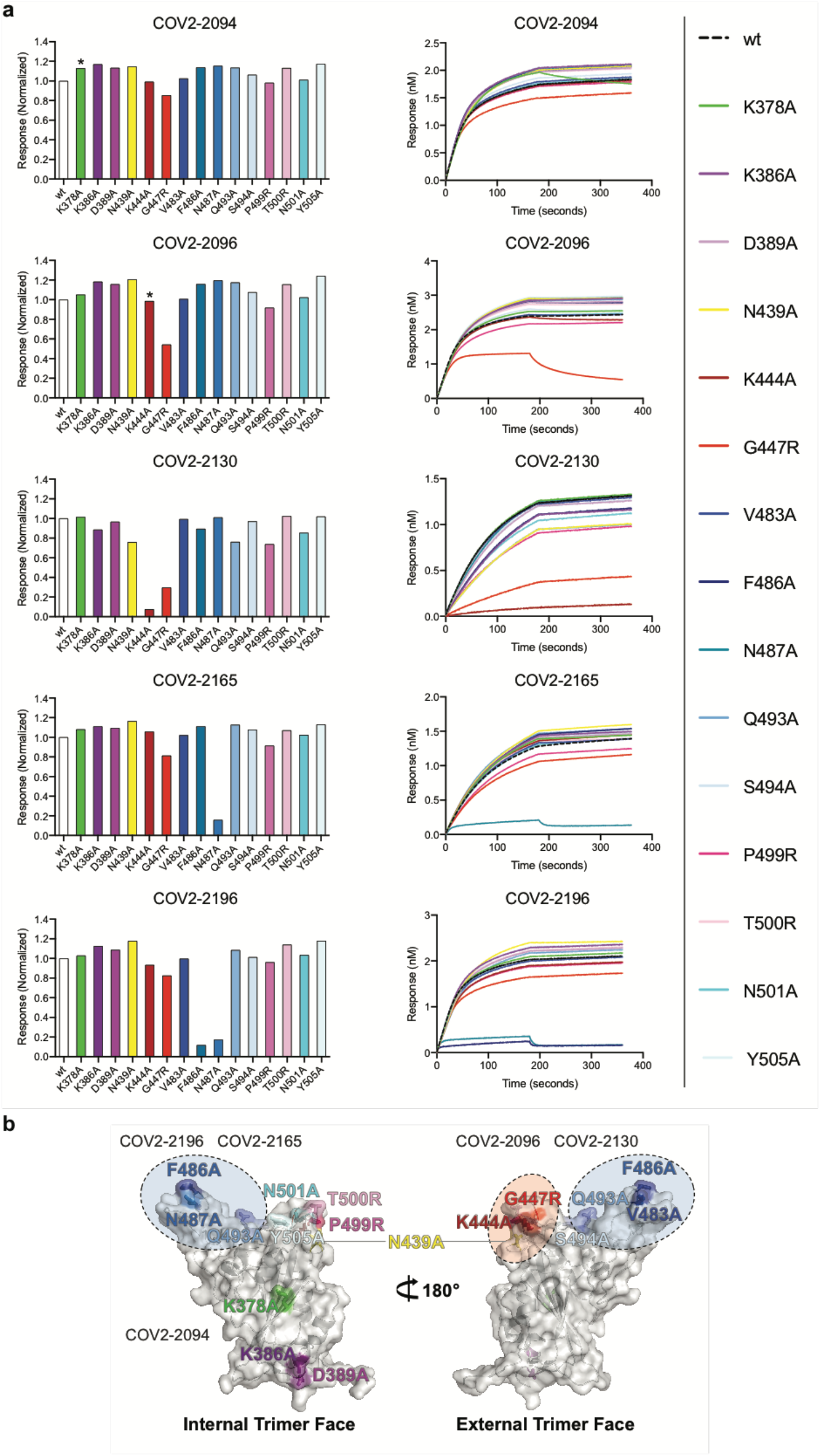
Mapping of mAb critical contact residues by alanine and arginine mutagenesis and biolayer interferometry. **a.** *Left*: Response values for mAb binding to *wt* or mutant S_RBD_ constructs normalized to *wt*. Asterisks denote residues where increased dissociation of mAb was observed, likely indicating the residue is proximal to mAb epitope. *Right*: full response curves for mAb association and dissociation with *wt* or mutant S_RBD_ constructs. **b.** Structure of the RBD highlighting the critical contact residues for several mAbs and their location on the structure.

**Extended Data Table 1.** Neutralization IC_50_, hACE2 blocking IC_50_, and EC_50_ values for binding to S2P_ecto_ or S_RBD_ antigens for mAb panel.

| **MAB** | **Neutralization**<br>**IC<sub>50</sub>, ng/mL** | **hACE2**<br>**blocking**<br>**IC<sub>50</sub>, ng/mL** | **S2P<sub>ecto</sub>**<br>**binding**<br>**EC<sub>50</sub>,**<br>**ng/mL** | **S<sub>RBD</sub>**<br>**binding**<br>**EC<sub>50</sub>,**<br>**ng/mL** | **SARS-CoV**<br>**S2P<sub>ecto</sub>**<br>**binding**<br>**EC<sub>50</sub>,**<br>**ng/mL** |
| --- | --- | --- | --- | --- | --- |
| COV2-2015 | 892 | 68 | 2.2 | 5.1 | 2.7 |
| COV2-2050 | 80 | 63 | 1.7 | 1.6 | — |
| COV2-2068 | 478 | 166 | 7.4 | 1.2 | — |
| COV2-2072 | 26 | 29 | 0.8 | 1.0 | — |
| COV2-2082 | 204 | 43 | 1.1 | 0.6 | 19.9 |
| COV2-2094 | 154 | 53 | 1.8 | 1.4 | 11.7 |
| COV2-2096 | 59 | 67 | 1.0 | 1.0 | — |
| COV2-2098 | 1,029 | 48 | 1.3 | 0.7 | — |
| COV2-2103 | 1,969 | 79 | 2.9 | 1.1 | — |
| COV2-2113 | 1,041 | 60 | 1.8 | 0.6 | — |
| COV2-2130 | 107 | 61 | 1.5 | 0.7 | — |
| COV2-2165 | 332 | 62 | 1.4 | 0.6 | — |
| COV2-2196 | 15 | 48 | 1.2 | 1.1 | — |
| COV2-2258 | 989 | 76 | 2.5 | 1.6 | 11.2 |
| COV2-2268 | 371 | 2,198 | 91.2 | 5.3 | — |
| COV2-2308 | 394 | 75 | 2.7 | 1.9 | — |
| COV2-2353 | 2,891 | 1,750 | 48.0 | 10.9 | — |
| COV2-2354 | 1,105 | 67 | 2.6 | 3.5 | — |
| COV2-2381 | 42 | 27 | 0.8 | 1.0 | — |
| COV2-2489 | 4,378 | — | 13.0 | — | — |
| COV2-2479 | 48 | 50 | 0.4 | 0.3 | — |
| COV2-2499 | 27 | 57 | 1.4 | 1.1 | — |
| COV2-2539 | 274 | 98 | 1.1 | 0.7 | — |
| COV2-2562 | 348 | 154 | 1.0 | 0.6 | — |
| COV2-2676 | 3,247 | — | 12.0 | — | — |
| COV2-2677 | 1,618 | 107 | 2.4 | 0.7 | — |
| COV2-2733 | 356 | 71 | 2.4 | 0.6 | — |
| COV2-2752 | 349 | 53 | 1.9 | 0.7 | — |
| COV2-2780 | 478 | 10,000 | 86.0 | 3.1 | — |
| COV2-2807 | 907 | 1,753 | 13.0 | 55.1 | — |
| COV2-2812 | 1,020 | 1,387 | 45.4 | 2.1 | — |
| COV2-2813 | 555 | 206 | 5.3 | 1.0 | — |
| COV2-2819 | 114 | 60 | 0.9 | 0.8 | — |
| COV2-2828 | 100 | 100 | 3.2 | 1.4 | 338 |
| COV2-2832 | 70 | 47 | 1.3 | 1.1 | — |
| COV2-2835 | 190 | 91 | 3.4 | 1.2 | — |
| COV2-2841 | 1,065 | 91 | 3.3 | 1.5 | — |
| COV2-2919 | 1,091 | 1,000 | 153.0 | 90.3 | — |
| COV2-2955 | 127 | 54 | 0.8 | 0.5 | — |
| COV2-3025 | 37 | 41 | 1.1 | 1.1 | — |
| rCR3022 | — | — | 10.2 | 1.1 | 5.2 |
| r2D22 | — | — | — | — | — |

— : indicates no detectable binding or neutralization at the highest tested concentration

