## Supplemental Methods for "Potently neutralizing human antibodies that block SARS-CoV-2 receptor binding and protect animals"

### Online Methods

**Antibodies.** The human antibodies studied in this paper were isolated from blood samples from two subjects in North America with previous laboratory-confirmed symptomatic SARS-CoV-2 infection that was acquired in China. The original clinical studies to obtain specimens after written informed consent were previously described[1] and had been approved by the Institutional Review Board of Vanderbilt University Medical Center and the Research Ethics Board of the University of Toronto. The subjects (a 56-year-old male and a 56-year-old female) are a married couple and residents of Wuhan, China who traveled to Toronto, Canada, where PBMCs were obtained by leukopheresis 50 days after symptom onset. The antibodies were isolated using diverse tools for isolation and cloning of single antigen-specific B cells and the antibody variable genes encoding monoclonal antibodies[1].

**Cell culture.** Vero E6 (CRL-1586, American Type Culture Collection (American Type Culture Collection, ATCC), Vero CCL81 (ATCC), HEK293 (ATCC), and HEK293T (ATCC) were maintained at 37°C in 5% $CO_2$ in Dulbecco's minimal essential medium (DMEM) containing 10% (vol/vol) heat-inactivated fetal bovine serum (FBS), 10 mM HEPES pH 7.3, 1 mM sodium pyruvate, 1× non-essential amino acids, and 100 U/mL of penicillin–streptomycin. Vero-furin cells were obtained from T. Pierson (NIH) and have been described previously[2]. Expi293F cells (ThermoFisher Scientific, A1452) were maintained at 37°C in 8% $CO_2$ in Expi293F Expression Medium (ThermoFisher Scientific, A1435102). ExpiCHO cells (ThermoFisher Scientific, A29127) were maintained at 37°C in 8% $CO_2$ in ExpiCHO Expression Medium (ThermoFisher Scientific, A2910002). Mycoplasma testing of Expi293F and ExpiCHO cultures was performed on a monthly basis using a PCR-based mycoplasma detection kit (ATCC, 30-1012K).

**Viruses.** SARS-CoV-2 strain 2019 n-CoV/USA_WA1/2020 was obtained from the Centers for Disease Control and Prevention (a gift from Natalie Thornburg). Virus was passaged in Vero CCL81 cells and titrated by plaque assay on Vero E6 cells. All work with infectious SARS-CoV-2 was approved by the Washington University School of Medicine or UNC-Chapel Hill Institutional Biosafety Committees and conducted in approved BSL3 facilities using appropriate powered air purifying respirators and personal protective equipment.

**Recombinant antigens and proteins.** A gene encoding the ectodomain of a prefusion conformation-stabilized SARS-CoV-2 spike ($S2P_{ecto}$) protein was synthesized and cloned into a DNA plasmid expression vector for mammalian cells. A similarly designed S protein antigen with two prolines and removal of the furin cleavage site for stabilization of the prefusion form of S was reported previously[3]. Briefly, this gene includes the ectodomain of SARS-CoV-2 (to residue 1,208), a T4 fibritin trimerization domain, an AviTag site-specific biotinylation sequence, and a C-terminal 8x-His tag. To stabilize the construct in the prefusion conformation, we included substitutions K986P and V987P and mutated the furin cleavage site at residues 682-685 from RRAR to ASVG. This recombinant spike 2P-stabilized protein (designated here as $S2P_{ecto}$) was isolated by metal affinity chromatography on HisTrap Excel columns (GE Healthcare), and protein preparations were purified further by size-exclusion chromatography on a Superose 6 Increase 10/300 column (GE Healthcare). The presence of trimeric, prefusion conformation S protein was verified by negative-stain electron microscopy[1]. For electron microscopy with S and Fabs, we expressed a variant of $S2P_{ecto}$ lacking an AviTag but containing a C-terminal Twin-Strep-tag, similar to that described previously[3]. Expressed protein was isolated by metal affinity chromatography on HisTrap Excel columns (GE Healthcare), followed by further purification on a StrepTrap HP column (GE Healthcare) and size-exclusion chromatography on

TSKgel G4000SW $_{XL}$ (TOSOH). To express the $S_{RBD}$ subdomain of SARS-CoV-2 S protein, residues 319-541 were cloned into a mammalian expression vector downstream of an IL-2 signal peptide and upstream of a thrombin cleavage site, an AviTag, and a 6x-His tag. RBD protein fused to mouse IgG1 Fc domain (designated RBD-mFc), was purchased from Sino Biological (40592-V05H). For epitope mapping by alanine scanning, SARS-CoV-2 RBD (residues 334-526) or RBD single mutation variants were cloned with an N-terminal CD33 leader sequence and C-terminal GSSG linker, AviTag, GSSG linker, and 8xHisTag. Spike proteins were expressed in FreeStyle 293 cells (Thermo Fisher) and isolated by affinity chromatography using a HisTrap column (GE Healthcare), followed by size exclusion chromatography with a Superdex200 column (GE Healthcare). Purified proteins were analyzed by SDS-PAGE to ensure purity and appropriate molecular weights.

**Electron microscopy (EM) stain grid preparation, imaging and processing of SARS-CoV-2 $S2P_{ecto}$ protein or $S2P_{ecto}$/Fab complexes.** To perform EM imaging, Fabs were produced by digesting recombinant chromatography-purified IgGs using resin-immobilized cysteine protease enzyme (FabALACTICA, Genovis). The digestion occurred in 100 mM sodium phosphate, 150 mM NaCl pH 7.2 for ~16 hrs at RT. In order to remove cleaved Fc and intact IgG, the digestion mix was incubated with CaptureSelect Fc resin (Genovis) for 30 min at RT in PBS buffer. If needed, the Fab was buffer exchanged into Tris buffer by centrifugation with a Zeba spin column (Thermo Scientific).

For screening and imaging of negatively-stained (NS) SARS-CoV-2 $S2P_{ecto}$ protein in complex with human Fabs, the proteins were incubated for ~1 hr and approximately 3 µL of the sample at concentrations of about 10 to 15 µg/mL was applied to a glow discharged grid with continuous carbon film on 400 square mesh copper EM grids (Electron Microscopy Sciences). The grids were stained with 0.75% uranyl formate (UF)[4]. Images were recorded on a Gatan US4000 4k × 4k CCD camera using an FEI TF20 (TFS) transmission electron microscope operated at 200 keV and control with SerialEM[5]. All images were taken at 50,000× magnification with a pixel size of 2.18 Å/pix in low-dose mode at a defocus of 1.5 to 1.8 µm. Total dose for the micrographs was ~25 to 38 $e^{-}$/Å$^{2}$. Image processing was performed using the cryoSPARC software package[6]. Images were imported, and particles were CTF estimated. The images then were denoised and picked with Topaz[7]. The particles were extracted with a box size of 256 pixels and binned to 128 pixels. 2D class averages were performed and good classes selected for *ab-initio* model and refinement without symmetry. For EM model docking of SARS-CoV-2 S2P$_{ecto}$ protein, the closed model (PDB: 6VXX) was used in Chimera[8] for docking to the EM map (see also Extended Data Table 2 for details). For the SARS-CoV-2 S2P$_{ecto}$/Fab COV2-2165 and SARS-CoV-2 S2P$_{ecto}$/Fab COV2-2165 complexes, the open model of SARS-CoV-2 (PDB: 6VYB) and Fab (Fab: 12E8) was used in Chimera for docking to the EM maps (see also Extended Data Table 2 for details). For the SARS-Cov-2 S2P$_{ecto}$/Fab COV2-2130 complex, the closed model and Fab (PDB: 12E8) were used in Chimera for docking to the EM map (see also Extended Data Table 2 for details). All images were made with Chimera.

**MAb production and purification.** Sequences of mAbs that had been synthesized (Twist Bioscience) and cloned into an IgG1 monocistronic expression vector (designated as pTwist-mCis_G1) were used for mammalian cell culture mAb secretion. This vector contains an enhanced 2A sequence and GSG linker that allows simultaneous expression of mAb heavy and light chain genes from a single construct upon transfection[9]. We previously described microscale expression of mAbs in 1 mL ExpiCHO cultures in 96-well plates[1]. For larger scale mAb expression, we performed transfection (1 to 300 mL per antibody) of CHO cell cultures using the Gibco™ ExpiCHO™ Expression System and protocol for 50 mL mini bioreactor tubes (Corning) as described by the vendor. Culture supernatants were purified using HiTrap MabSelect SuRe (Cytiva, formerly GE Healthcare Life Sciences) on a 24-column parallel protein chromatography system (Protein BioSolutions). Purified mAbs were buffer-exchanged into PBS, concentrated using Amicon® Ultra-4 50KDa Centrifugal Filter Units (Millipore Sigma) and stored at 4°C until use.

**ELISA binding assays.** Wells of 96-well microtiter plates were coated with purified recombinant SARS-CoV-2 S protein or SARS-CoV-2 $S_{RBD}$ protein at 4°C overnight. Plates were blocked with 2% non-fat dry milk and 2% normal goat serum in DPBS containing 0.05% Tween-20 (DPBS-T) for 1 hr. The bound antibodies were detected using goat anti-human IgG conjugated with HRP (horseradish peroxidase) (Southern Biotech) and TMB (3,3′,5,5′-tetramethylbenzidine) substrate (Thermo Fisher Scientific). Color development was monitored, 1N hydrochloric acid was added to stop the reaction, and the absorbance was measured at 450 nm using a spectrophotometer (Biotek). For dose-response assays, serial dilutions of purified mAbs were applied to the wells in triplicate, and mAb binding was detected as detailed above. Half-maximal effective concentration ($EC_{50}$) values for binding were determined using Prism v8.0 software (GraphPad) after log transformation of mAb concentration using sigmoidal dose-response nonlinear regression analysis.

**RBD minimal hACE2 recognition motif peptide binding ELISA.** Wells of 384-well microtiter plates were coated with 1 µg/mL streptavidin at 4°C overnight. Plates were blocked with 0.5% BSA in DPBS containing 0.05% Tween-20 (DPBS-T) for 1 hr. Plates were washed 4x with 1x PBST and 2 μg/mL biotinylated-ACE2 binding motif peptide (cat. # LT5578, from LifeTein, LLC) was added to bind streptavidin for 1 hr at RT. Purified mAbs were diluted in blocking buffer, added to the wells, and incubated for 1 hr at RT. The bound antibodies were detected using goat anti-human IgG conjugated with HRP (horseradish peroxidase) (cat. # 2014-05, Southern Biotech) and TMB (3,3′,5,5′-tetramethylbenzidine) substrate (ThermoFisher Scientific). Color development was monitored, 1N hydrochloric acid was added to stop the reaction, and the absorbance was measured at 450 nm using a spectrophotometer (Biotek). For dose-response assays, serial 3-fold dilutions starting at 10 μg/mL concentration of purified mAbs were applied to the wells in triplicate, and mAb binding was detected as detailed above.

**Analysis of binding of antibodies to variant RBD proteins with alanine or arginine point mutations.** Biolayer light interferometry (BLI) was performed using an Octet RED96 instrument (ForteBio; Pall Life Sciences) and wild-type RBD protein or a mutant RBD protein with a single amino acid change at defined positions to alanine or arginine. Binding of the RBD proteins were confirmed by first capturing octa-His-tagged RBD wild-type or mutant protein from a 10 μg/mL (≈200 nM) solution onto Penta-His biosensors for 300 sec. The biosensor tips then were submerged in binding buffer (PBS/0.2% Tween 20) for a 60 sec wash, followed by immersion in a solution containing 150 nM of mAb for 180 sec (association), followed by a subsequent immersion in binding buffer for 180 sec (dissociation). Response for each RBD mutant protein was normalized to that of the wild-type RBD protein.

**Focus reduction neutralization test (FRNT).** Serial dilutions of mAbs were incubated with $10^2$ FFU of SARS-CoV-2 for 1 hr at 37°C. The mAb–virus complexes were added to Vero E6 cell culture monolayers in 96-well plates for 1 hr at 37°C. Subsequently, cells were overlaid with 1% (w/v) methylcellulose in Minimum Essential Medium (MEM) supplemented to contain 2% heat-inactivated FBS. Plates were fixed 30 hrs later by removing overlays and fixed with 4% PFA in PBS for 20 min at room temperature. The plates were incubated sequentially with 1 μg/mL of rCR3022 anti-S antibody[10] and horseradish-peroxidase (HRP)-conjugated goat anti-human IgG in PBS supplemented with 0.1% (w/v) saponin (Sigma) and 0.1% bovine serum albumin (BSA). SARS-CoV-2-infected cell foci were visualized using TrueBlue peroxidase substrate (KPL) and quantitated on an ImmunoSpot 5.0.37 Macro Analyzer (Cellular Technologies). Data were processed using Prism software version 8.0 (GraphPad). $IC_{50}$ values were determined by nonlinear regression analysis using the Prism software.

**Generation of S protein pseudotyped lentivirus.** Suspension 293 cells were seeded and transfected with a third-generation HIV-based lentiviral vector expressing luciferase along with packaging plasmids encoding for the following:  SARS-CoV-2 spike protein with a C-terminal 19 amino acid deletion, Rev, and Gag-pol. Medium was changed 16 to 20 hrs after transfection, and the supernatant containing virus was harvested 24 hrs later. Cell debris was removed by low-speed centrifugation, and the supernatant was passed through a 0.45 μm filter unit. The pseudovirus was pelleted by ultracentrifugation and resuspended in PBS for a 100-fold concentrated stock.

**Pseudovirus neutralization assay.** Serial dilutions of mAbs were prepared in a 384-well microtiter plate and pre-incubated with pseudovirus for 30 minutes at 37°C, to which 293 cells that stably express human ACE2 were added. The plate was returned to the 37°C incubator, and then 48 hrs later luciferase activity measured on an EnVision 2105 Multimode Plate Reader (Perkin Elmer) using the Bright-Glo™

Luciferase Assay System (Promega), according to manufacturer's recommendations. Percent inhibition was calculated relative to pseudovirus-alone control. $IC_{50}$ values were determined by nonlinear regression using the Prism software version 8.1.0 (GraphPad). The average $IC_{50}$ value for each antibody was determined from a minimum of 3 independent experiments.

**Measurement of synergistic neutralization by an antibody combination.** Synergy was defined as higher neutralizing activity mediated by a cocktail of two mAbs when compared to that mediated by individual mAbs at the same total concentration of antibodies *in vitro*. To assess if two mAbs synergize in a cocktail to neutralize SARS-CoV-2, we used a previously reported approach to quantitate synergy[11]. To evaluate the significance of the beneficial effect from combining mAbs, the observed combination responses (dose-response matrix) were compared with the expected responses calculated by means of synergy scoring models[11]. Virus neutralization was measured in a conventional focus reduction neutralization test (FRNT) assay using wild-type SARS-CoV-2 and Vero E6 cell culture monolayers. Individual mAbs COV2-2196 and COV2-2130 were mixed at different concentrations to assess neutralizing activity of different ratios of mAbs in the cocktail. Specifically, each of seven-fold dilutions of mAb COV2-2130 (starting from 500 ng/mL) was mixed with each of the nine dilutions of mAb COV2-2196 (starting from 500 ng/mL) in a total volume of 50 μL of per each condition and then incubated with 50 μL of live SARS-CoV-2 in cell culture medium (RPMI-1640 medium supplemented with 2% FBS) before applying to confluent Vero E6 cells grown in 96-well plates. The control values included those for determining dose-response of the neutralizing activity measured separately for the individual mAb COV2-2196 or COV2-2130, which were assessed at the same doses as in the cocktail. Each measurement was performed in duplicate. We next calculated percent virus neutralization for each condition and then calculated the synergy score value, which defined interaction between these two mAbs in the cocktail. A synergy score of less than -10 indicates antagonism, a score from -10 to 10 indicates an additive effect, and a score greater than 10 indicates a synergistic effect[11].

**MAb quantification.** Quantification of purified mAbs was performed by UV spectrophotometry using a NanoDrop spectrophotometer and accounting for the extinction coefficient of human IgG.

**Competition-binding analysis through biolayer interferometry.** Anti-mouse IgG Fc capture biosensors (FortéBio 18-5089) on an Octet HTX biolayer interferometry instrument (FortéBio) were soaked for 10 minutes in 1x kinetics buffer (Molecular Devices 18-1105), followed by a baseline signal measurement for 60 seconds. Recombinant SARS-CoV-2 RBD fused to mouse IgG1 (RBD-mFc, Sino Biological 40592-V05H) was immobilized onto the biosensor tips for 180 seconds. After a wash step in 1x kinetics buffer for 30 seconds, the reference antibody (5 µg/mL) was incubated with the antigen-containing biosensor for 600 seconds. Reference antibodies included the SARS-CoV human mAb CR3022 and COV2-2196. After a wash step in 1x kinetics buffer for 30 seconds, the biosensor tips then were immersed into the second antibody (5 µg/mL) for 300 seconds. Maximal binding of each antibody was normalized to a buffer-only control. Self-to-self blocking was subtracted. Comparison between the maximal signal of each antibody was used to determine the percent binding of each antibody. A reduction in maximum signal to <33% of un-competed signal was considered full competition of binding for the second antibody in the presence of the reference antibody. A reduction in maximum signal to between 33 to 67% of un-competed was considered intermediate competition of binding for the second antibody in the presence of the reference antibody. Percent binding of the maximum signal >67% was considered absence of competition of binding for the second antibody in the presence of the reference antibody.

**hACE-2 inhibition analysis.** Wells of 384-well microtiter plates were coated with purified recombinant SARS-CoV-2 $S2P_{ecto}$ protein at 4°C overnight. Plates were blocked with 2% non-fat dry milk and 2% normal goat serum in DPBS-T for 1 hr. For screening assays, purified mAbs from microscale expression were diluted two-fold in blocking buffer starting from 10 µg/mL in triplicate, added to the wells (20 µL/well), and incubated for 1 hr at ambient temperature. Recombinant human hACE2 with a C-terminal FLAG tag protein was added to wells at 2 µg/mL in a 5 µL/well volume (final 0.4 µg/mL concentration of ACE2) without washing of antibody and then incubated for 40 min at ambient temperature. Plates were washed, and bound ACE2 was detected using HRP-conjugated anti-FLAG antibody (Sigma) and TMB substrate. ACE2 binding without antibody served as a control. The signal obtained for binding of the ACE2 in the presence of each dilution of tested antibody was expressed as a percentage of the ACE2 binding without antibody after subtracting the background signal. For dose-response assays, serial dilutions of purified mAbs were applied to the wells in triplicate, and mAb binding was detected as detailed above. Half-maximal inhibitory concentration ($IC_{50}$) values for inhibition by mAb of $S2P_{ecto}$ protein binding to ACE2 was determined after log transformation of antibody concentration using sigmoidal dose-response nonlinear regression analysis (Prism software, GraphPad Prism version 8.0).

**ACE2 blocking assay using biolayer interferometry biosensor.** Anti-mouse IgG biosensors on an Octet HTX biolayer interferometry instrument (ForteBio) were soaked for 10 minutes in 1x kinetics buffer, followed by a baseline signal measurement for 60 seconds. Recombinant SARS-CoV-2 RBD fused to mouse IgG1 (RBD-mFc, Sino Biological 40592-V05H) was immobilized onto the biosensor tips for 180 seconds. After a wash step in 1x kinetics buffer for 30 seconds, the antibody (5 μg/mL) was incubated with the antigen-coated biosensor for 600 seconds. After a wash step in 1x kinetics buffer for 30 seconds, the biosensor tips then were immersed into the ACE2 receptor (20 μg/mL) (Sigma-Aldrich SAE0064) for 300 seconds. Maximal binding of ACE2 was normalized to a buffer-only control. Percent binding of ACE2 in the presence of antibody was compared to ACE2 maximal binding. A reduction in maximal signal to <30% was considered ACE2 blocking.

**High-throughput competition-binding analysis.** Wells of 384-well microtiter plates were coated with purified recombinant SARS-CoV-2 S protein at 4°C overnight. Plates were blocked with 2% BSA in DPBS containing 0.05% Tween-20 (DPBS-T) for 1 hr. Micro-scale purified unlabeled mAbs were diluted ten-fold in blocking buffer, added to the wells (20 μL/well) in quadruplicates, and incubated for 1 hr at ambient temperature. A biotinylated preparation of a recombinant mAb based on the variable gene sequence of the previously described mAb CR3022[12] and also newly identified mAbs COV2-2096, -2130, and -2196 that recognized distinct antigenic regions of the SARS-CoV-2 S protein were added to each of four wells with the respective mAb at 2.5 μg/mL in a 5 μL/well volume (final 0.5 μg/mL concentration of biotinylated mAb) without washing of unlabeled antibody and then incubated for 1 hr at ambient temperature. Plates were washed, and bound antibodies were detected using HRP-conjugated avidin (Sigma) and TMB substrate. The signal obtained for binding of the biotin-labeled reference antibody in the presence of the unlabeled tested antibody was expressed as a percentage of the binding of the reference antibody alone after subtracting the background signal. Tested mAbs were considered competing if their presence reduced the reference antibody binding to less than 41% of its maximal binding and non-competing if the signal was greater than 71%. A level of 40–70% was considered intermediate competition.

**Binding analysis of mAbs to alanine or arginine RBD mutants.** Biolayer light interferometry was performed using an Octet RED96 instrument (ForteBio; Pall Life Sciences). Binding was confirmed by first capturing octa-His-tagged RBD mutants 10 μg/mL (≈200 nM) onto Penta-His biosensors for 300 s. The biosensors then were submerged in binding buffer (PBS/0.2% TWEEN 20) for a wash for 60 sec followed by immersion in a solution containing 150 nM of mAbs for 180 sec (association), followed by a subsequent immersion in binding buffer for 180 sec (dissociation). Response for each RBD mutant was normalized to that of wild-type RBD.

**Mouse experiments using human *hACE2*-transduced mice.** Animal studies were carried out in accordance with the recommendations in the Guide for the Care and Use of Laboratory Animals of the National Institutes of Health. The protocols were approved by the Institutional Animal Care and Use Committee at the Washington University School of Medicine (assurance number A3381–01). Virus inoculations were performed under anesthesia that was induced and maintained with ketamine hydrochloride and xylazine, and all efforts were made to minimize animal suffering.

BALB/c mice were purchased from Jackson Laboratories (strain 000651). Female mice (10-11-week-old) were given a single intraperitoneal injection of 2 mg of anti-Ifnar1 mAb (MAR1-5A3 [13], Leinco) one day before intranasal administration of 2.5 x $10^8$ PFU of AdV-hACE2. Five days after AdV transduction, mice were inoculated with 4 x $10^5$ PFU of SARS-CoV-2 by the intranasal route.

Anti-SARS-CoV-2 human mAbs or isotype control mAbs were administered 24 hours prior to SARS-CoV-2 inoculation. Weights were monitored on a daily basis, and animals were sacrificed at days 5 or 7 post-infection, and tissues were harvested.

**Measurement of viral burden.** Tissues were weighed and homogenized with zirconia beads in a MagNA Lyser instrument (Roche Life Science) in 1 ml of DMEM media supplemented with 2% heat-inactivated FBS. Tissue homogenates were clarified by centrifugation at 10,000 rpm for 5 min and stored at −80°C. RNA was extracted using MagMax mirVana Total RNA isolation kit (Thermo Scientific) and a Kingfisher Flex 96 well extraction machine (Thermo Scientific). TaqMan primers were designed to target a conserved region of the N gene using SARS-CoV-2 (MN908947) sequence as a guide (L Primer: ATGCTGCAATCGTGCTACAA; R primer: GACTGCCGCCTCTGCTC; probe: /56-FAM/TCAAGGAAC/ZEN/AACATTGCCAA/3IABkFQ/). To establish an RNA standard curve, we generated concatenated segments of the N gene in a gBlocks fragment (IDT) and cloned this into the PCR-II topo vector (Invitrogen). The vector was linearized, and *in vitro* T7-DNA-dependent RNA transcription was performed to generate materials for a quantitative standard curve.

**Cytokine and chemokine mRNA measurements**. RNA was isolated from lung homogenates at 7 dpi as described above. cDNA was synthesized from DNase-treated RNA using the High-Capacity cDNA Reverse Transcription kit (Thermo Scientific) with the addition of RNase inhibitor, following the manufacturer's protocol. Cytokine and chemokine expression was determined using TaqMan Fast Universal PCR master mix (Thermo Scientific) with commercial primers/probe sets specific for *IFNγ* (IDT: Mm.PT.58.41769240), *IL*

*6* (Mm.PT.58.10005566), *CXCL10* (Mm.PT.58.43575827), *CCL2* (Mm.PT.58.42151692 and results were normalized to *GAPDH* (Mm.PT.39a.1) levels. Fold change was determined using the $2^{-\Delta\Delta Ct}$ method comparing anti-SARS-CoV-2 specific or isotype control mAb-treated mice to naïve controls.

**Mouse experiments using wild-type mice.** Animal studies were carried out in accordance with the recommendations in the Guide for the Care and Use of Laboratory Animals of the National Institutes of Health. The protocols were approved by the Institutional Animal Care and Use Committee at the UNC Chapel Hill School of Medicine (NIH/PHS Animal Welfare Assurance Number is D16-00256 (A3410-01)). Virus inoculations were performed under anesthesia that was induced and maintained with ketamine hydrochloride and xylazine, and all efforts were made to minimize animal suffering.

**Mouse adapted SARS-CoV-2 (MA-SARS-CoV-2) virus.** The virus was generated as described previously[14]. Virus was propagated in Vero E6 cells grown in DMEM with 10% Fetal Clone II and 1% Pen/Strep. Virus titer was determined by plaque assay. Briefly, virus was serial diluted and inoculated onto confluent monolayers of Vero E6 cells, followed by agarose overlay. Plaques were visualized on day 2 post-infection after staining with neutral red dye.

**Wild-type mice**. 12-month-old BALB/c mice from Envigo were used in experiments. Mice were acclimated in the BSL3 for at least 72 hrs prior to start of experiments. At 6 hrs prior to infection, mice were prophylactically treated with 200 µg of human monoclonal antibodies via intraperitoneal injection. The next day, mice were anesthetized with a mixture of ketamine and xylazine and intranasally infected with $10^5$ PFU of MA-SARS-CoV-2 diluted in PBS. Daily weight loss was measured, and at two days post-infection mice were euthanized by isoflurane overdose prior to tissue harvest.

**Plaque assay of lung tissue homogenates**. The lower lobe of the right lung was homogenized in 1 mL PBS using a MagnaLyser (Roche). Serial dilutions of virus were titered on Vero E6 cell culture monolayers, and virus plaques were visualized by neutral red staining at two days after inoculation. The limit of detection for the assay is 100 PFU per lung.

**Quantification and statistical analysis.** The descriptive statistics mean ± SEM or mean ± SD were determined for continuous variables as noted. Technical and biological replicates are described in the figure legends. In the mouse studies, the comparison of weight change curves was performed using repeated measurements two-way ANOVA with Tukey's post-test using Prism v8.0 (GraphPad). Viral burden and gene expression measurements were compared using Kruskal-Wallis ANOVA with Dunn’s post-test using Prism v8.0 (GraphPad). Synergy score and dose-response matrix analysis were performed using a web application SynergyFinder[11].

**References for Online Methods**

1 Zost SJ, G. P., Chen RE, Case JB, Reidy JX, Trivette A, Nargi RS, Sutton RE, Suryadevara N, Chen EC, Binshtein E, Shrihari S, Ostrowski M, Chu HY, Didier JE, MacRenaris KW, Jones T, Day S, Myers L, Lee FE-H, Nguyen DC, Sanz I, Martinez DR, Baric RS, Thackray LB,. Diamond MS, Carnahan RH, Crowe JE Jr. Rapid isolation

and profiling of a diverse panel of human monoclonal antibodies targeting the SARS-CoV-2 spike protein. *bioRxiv* 2020.05.12.091462; doi: https://doi.org/10.1101/2020.05.12.091462 (2020)

2 Mukherjee, S. *et al.* Enhancing dengue virus maturation using a stable furin over-expressing cell line. *Virology* **497**, 33-40, doi:10.1016/j.virol.2016.06.022 (2016).

3 Wrapp, D. *et al.* Cryo-EM structure of the 2019-nCoV spike in the prefusion conformation. *Science* **367**, 1260-1263, doi:10.1126/science.abb2507 (2020).

4 Ohi, M., Li, Y., Cheng, Y. & Walz, T. Negative staining and image classification - Powerful tools in modern electron microscopy. *Biol. Proced. Online* **6**, 23-34, doi:10.1251/bpo70 (2004).

5 Mastronarde, D. N. Automated electron microscope tomography using robust prediction of specimen movements. *J. Struct. Biol.* **152**, 36-51, doi:10.1016/j.jsb.2005.07.007 (2005).

6 Punjani, A., Rubinstein, J. L., Fleet, D. J. & Brubaker, M. A. cryoSPARC: algorithms for rapid unsupervised cryo-EM structure determination. *Nat. Methods* **14**, 290-296, doi:10.1038/nmeth.4169 (2017).

7 Bepler, T., Noble, A. J., and Berger, B. Topaz-Denoise: general deep denoising models for cryoEM. *bioRxiv.* doi:10.1101/838920 (2019).

8 Pettersen, E. F. *et al.* UCSF Chimera--a visualization system for exploratory research and analysis. *J. Comput. Chem.* **25**, 1605-1612, doi:10.1002/jcc.20084 (2004).

9 Chng, J. *et al.* Cleavage efficient 2A peptides for high level monoclonal antibody expression in CHO cells. *MAbs* **7**, 403-412, doi:10.1080/19420862.2015.1008351 (2015).

10 Yuan, M. *et al.* A highly conserved cryptic epitope in the receptor binding domains of SARS-CoV-2 and SARS-CoV. *Science* **368**, 630-633, doi:10.1126/science.abb7269 (2020).

11 Ianevski, A., He, L., Aittokallio, T. & Tang, J. SynergyFinder: a web application for analyzing drug combination dose-response matrix data. *Bioinformatics* **33**, 2413-2415, doi:10.1093/bioinformatics/btx162 (2017).

12 ter Meulen, J. *et al.* Human monoclonal antibody as prophylaxis for SARS coronavirus infection in ferrets. *Lancet* **363**, 2139-2141, doi:10.1016/S0140-6736(04)16506-9 (2004).

13 Sheehan, K. C. *et al.* Blocking monoclonal antibodies specific for mouse IFN-alpha/beta receptor subunit 1 (IFNAR-1) from mice immunized by in vivo hydrodynamic transfection. *J. Interferon Cytokine Res* **26**, 804-819 (2006).

14 Dinnon KH, *et al.* A mouse-adapted SARS-CoV-2 model for the evaluation of COVID-19 medical countermeasures. *bioRxiv* 2020.05.06.081497; doi: https://doi.org/10.1101/2020.05.06.081497 (2020).
